# Network mechanisms in rapid-onset dystonia-parkinsonism

**DOI:** 10.1101/2024.10.14.618133

**Authors:** Meret Möller, Johanna A. Nieweler, Vadim V. Nikulin, Christoph van Riesen

## Abstract

**Background:** Rapid-onset dystonia-parkinsonism (RDP) is a rare neurological disorder caused by mutations in the ATP1A3 gene. Symptoms are characterized by a dystonia-parkinsonism. Recently, experimental studies have shown that the pathophysiology of the disease is based on a combined dysfunction of the cerebellum (CB) and basal ganglia (BG) and that blocking their interaction can alleviate the symptoms. The underlying network mechanisms have not been studied so far.

**Objective:** Our aim was to characterize neuronal network activity in the BG and CB and motor cortex in the ouabain model of RDP by site-specific infusion of ouabain.

**Methods:** Rats were chronically infused with ouabain either in the CB, striatum (STR) or at both places simultaneously. Motor behavior was scored using published rating systems. Parallel in vivo recordings of local field potentials (LFP) from M1, deep cerebellar nuclei (DCN) and substantia nigra reticulata (SNr) were performed. Data were compared to untreated controls.

**Results:** Ouabain infusion into the cerebellum produced severe dystonia that was associated with increased high-frequency gamma oscillations in the DCNs, which were subsequently transmitted to the BG and M1. Striatal infusion led to parkinsonism and elevated beta-oscillations in SNr that were transmitted to the CB and M1. The simultaneous application of STRs and CB with ouabain resulted in dystonia-parkinsonism and increased beta oscillations in BG, CB, and M1.

**Conclusion:** We demonstrate that symptom-specific beta and gamma oscillations can be transmitted between the BG and CB, which is likely to be very important for the understanding of disease mechanisms.

## Introduction

Rapid-onset dystonia-parkinsonism (RDP, DYT 12) is a rare neurological disorder that is characterized by abrupt onset of a combination of parkinsonism and dystonia. The disease is triggered by stressors, such as severe infections or physical exertion. After that, the disease remains stable, even when the stressors are no longer present. There is currently no effective therapy for this condition. Treatments used for disorders with a similar spectrum of symptoms (e.g. Parkinson’s disease and dystonia), including drug therapies and deep brain stimulation, are not effective.^1, 2^

RDP is caused by an autosomal dominant mutation in the ATP1A3 gene, which encodes the alpha-3 subunit isoform of the heterodimeric Na^+^-K^+^-ATPase.^3^ Brain Na^+^-K^+^-ATPases are essential for maintaining the resting membrane potential of neurons.^4^ They consist of a catalytic alpha subunit containing the Na^+^-K^+^-antiport and a regulatory beta subunit. The alpha 3 subunit is highly expressed in the basal ganglia (BG), thalamus, and cerebellum (CB).^5^ The missense mutations in the alpha-3 subunit identified in RDP result in a diminished capacity of neurons to perform their functions in situations of elevated metabolic demand.^6^ Given the essential role of ion homeostasis in various neuronal transport mechanisms (e.g., neurotransmitter transport), stress can precipitate a profound disruption in neuronal function, which in turn gives rise to the debilitating disease symptoms.^7^

The recently published ouabain animal model of RDP provides evidence for the simultaneous involvement of the basal ganglia (BG) and cerebellum (CB) in the pathogenesis of RDP.^8–10^ ouabain is a toxin that selectively blocks the alpha 3 subunit of the Na^+^-K^+^-ATPase. ^3^ It has been repeatedly shown that site-specific intracerebral infusion of ouabain can be employed to reliably model the diverse symptoms of RDP.^8, 10, 11^ Infusion of ouabain into the BG results in parkinsonism, whereas infusion into the CB gives rise to generalized dystonia. Combined infusion into the BG and CB leads to dystonia and parkinsonism, which is similar to the symptomatology of patients with RDP.^8, 10, 11^ It is noteworthy that after the induction of dystonia by ouabain infusion into the CB, blockade of the subcortical disynaptic connection between the deep cerebellar nuclei (DCN) and the striatum (STR) via the centrolateral thalamus (cl-thalamus) has been observed to alleviate symptoms.^8^ This evidence substantiates the hypothesis that the pathophysiology of RDP involves the basal ganglia and the cerebellum, and that it might be characterised by a dysfunctional subcortical interaction between the two motor control loops.

For a long time it was assumed that the BG and the CB were anatomically seperate, with the primary interaction between these loops occurring at the cortical level through slow multisynaptic pathways.^12–15^ However, recent cumulative evidence has highlighted the significance of the subcortical connection between the two in healthy and diseased states such as PD. ^16^

t has been demonstrated that there is a rapid bidirectional communication between the BG and CB via a subcortical disynaptic pathway. One of the pathways has its origin in the DCN and traverses through the cl-thalamus to the striatum. The other pathway commences at the subthalamic nucleus and proceeds via the pontine nuclei (pN) to the cerebellar cortex.^13, 17–20^ However, the present understanding of the bidirectional interaction between BG and CB in disease is limited. In particular, the underlying mechanisms through which this interaction occurs remain poorly understood.

For this reason, we conducted an investigation of the network mechanisms in the RDP- ouabain rat model through the use of parallel in vivo electrophysiological recordings of local field potentials (LFPs) from the basal ganglia, cerebellum, and M1, as well as the connecting network nodes. We chose to focus on LFPs because, unlike single unit activity, LFPs represent the cumulative activity of larger volumes of brain tissue, and because LFP analysis techniques allow a comprehensive examination of the interactions between distant brain areas.^21, 22^ Additionally, the translational significance of LFPs for developing and optimizing neuromodulatory therapies such as deep brain stimulation has been consistently demonstrated in the context of diverse movement disorders, including PD and dystonia.

In our study, we provide a detailed account of the changes in neuronal network activity observed in the BG and CB and between the them when either dystonia, parkinsonism, or dystonia-parkinsonism were induced by site-specific ouabain infusion. The results demonstrate that cerebellar infusion of ouabain resulted in the induction of generalized dystonia, which was associated with the occurrence of cerebellar high-frequency narrow-bandwidth gamma oscillations that were transmitted to the basal ganglia and M1. The infusion of ouabain into the striatum resulted in the generation of synchronized beta activity, which subsequently spread to the CB and M1. The combined infusion of ouabain into the STR and CB induced dystonia-parkinsonism with elevated beta oscillations in both motor control loops and M1. In conclusion, our findings represent the first demonstration of the basic mechanisms underlying the complex pathophysiological interaction of the basal ganglia and cerebellar motor control loops with M1 in RDP at a network level.

## Material and Methods

### 1. Material and methods

#### 1.1 Animals and materials

All experimental procedures were performed on male Wistar rats (n = 40, 310–495g, Charles River and ZTE Göttingen, Germany) in accordance with the European Union Directive 2010/63/EU and the German Animal Welfare Act (revised December 2018). Experiments were approved by the local animal welfare authority, and complied with local departmental and international guidelines. Animals were housed under standard conditions with a 12 h light/dark cycle and free access to food and water.

### 1.2 Experimental design

The objective of this study was to characterize the activity of the motor cortex, basal ganglia and cerebellar network in a rat model of RDP. Five different groups of animals were analyzed (n=8 for each group, Fig. 1). Based on recent publications ouabain was chronically infused either into the midline of the cerebellum (to induce dystonia; CB-group) or into the striatum bilaterally (to induce parkinsonism; STR-group) or into the cerebellum and striatum simultaneously (to induce dystonia-parkinsonism; STRCB-group). To account for the networks involved, overlapping and partially different recording sites were used for the CB- and STR-groups, which were therefore compared to two separate groups of untreated controls (see Figure 1 for an overview of recording sites). As only a reduced number of sites could be recorded in the STRCB-group due to spatial constraints, it was compared to a combination of both control groups.

**Fig.1:**
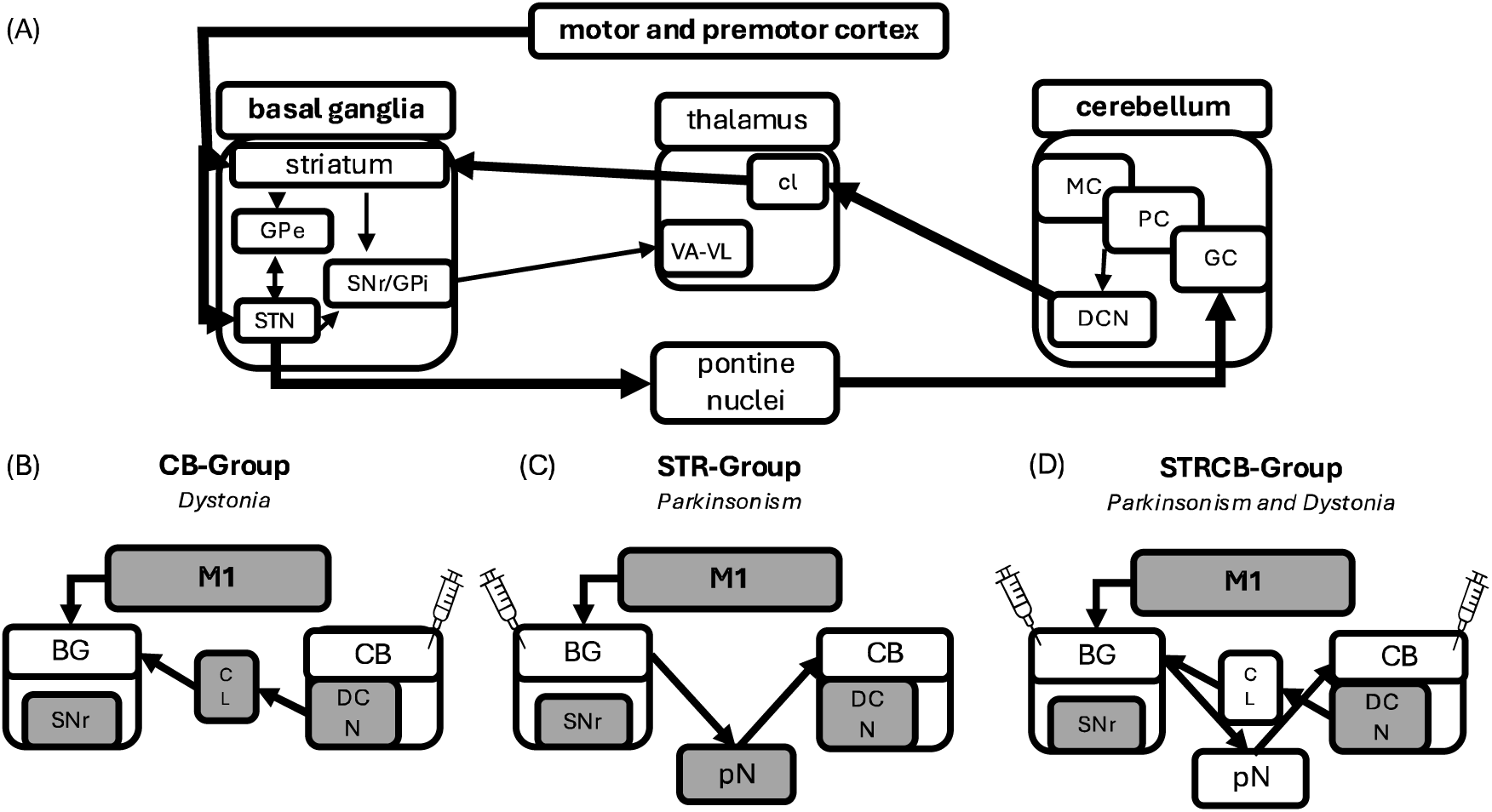
Schematic representation of the basal ganglia and cerebellar networks and their connections to the motor cortex. **(A)** Detailed overview. Inhibitory pathways shown with thin lines and excitatory pathways with thick lines. **(B, C, D)** Overview of the different experimental groups and the electrophysiological recording sites. Recording site are highlighted in grey. BG, basal ganglia. CB, cerebellum. Cl, centrolateral thalamus. GC, granule cell layer. GPe, external part of globus pallidus. M1, primary motor cortex. MC, molecular cell layer. PC, Purkinje cell layer. pN, pontine nuclei. STN, subthalamic nucleus. VA, ventroanterior thalamus. VL, ventrolateral thalamus.

### 1.3 Implantation of osmotic pumps

Osmotic pumps (Alzet, DURECT Corporation, CA, USA) were implanted stereotactically for chronic perfusion of the cerebellum and/or the striatum with ouabain. After induction and maintenance of isoflurane anesthesia, ophthalmic ointment (Dexpanthenol, Bepanthen ®, Bayer, Germany) was applied to prevent corneal dehydration. The animal was placed on a self-adjusting heating pad (37 ± 0.5°C) and transferred to a stereotaxic frame (David Kopf Instruments, Tujunga CA). The head was fixed in the flat-skull position using atraumatic ear bars. All stereotaxic coordinates were measured relative to bregma using a standard rat stereotaxic atlas.^23^ All stereotaxic coordinates for all experiments can be found in Suppl. Table 1 and 2. The cannulas were inserted into the midline of the CB and/or the STR bilaterally. Pumps were placed subcutaneously behind the neck. Carprofen (5 mg/kg, s.c., Pfizer, Germany) was administered as pain medication.

### 1.4 Behavioral assessment

Dystonia and parkinsonism were assessed using established scores (Locomotion Dysabililty Score (LDS), Dystonia Rating Scale, DRS)^8^ and documented by video recordings.

### 1.5 Electrophysiological recordings

Electrophysiological experiments were started after 72 h of chronic perfusion or when the animals reached a maximum score on one of the symptom scales. Anesthesia was induced and maintained with urethane (1.3 g/kg, i.p.). Electrophysiological recordings were performed as recently described in detail. ^24–27^ Stereotaxic coordinates can be found in Suppl. tabl. 1. Briefly, custom-made Ag/AgCl-electrodes (8 kΩ) were placed epidurally over the left primary motor cortex to record the electrocorticogram and over the contralateral cerebellum as a common reference. Additional Ag/AgCl-reference electrodes were placed in the epidural space near the microelectrodes as an alternative local reference. Two tungsten microelectrodes (1.2-1.5 MΩ; Science Products GmbH, Germany) were implanted in the substantia nigra pars reticulata and the dentate nucleus. An additional pair of tungsten electrodes was placed in the centrolateral-thalamus in the CB-group and in the pontine nuclei. in the STR-group. All microelectrodes were inserted under constant monitoring of multiunit activity to ensure optimal placement. All electrodes were fixed with dental cement. Electrophysiological recordings were done in a Faraday cage using a programmable neural data acquisition system (Omniplex, Plexon, TX, USA) with a 50Hz notch filter. The recorded wide-band signal (0.05–8000Hz; 40 kHz sampling rate) was bandpass-filtered and downsampled (0.05-250 Hz; 1 kHz) for later offline analysis of LFP data.

### 1.6 Electrophysiological data analysis

LFP data were analyzed using Spike2 (v8.02e, Cambridge Electronic Design, UK) and MatLab (2022, MathWorks, MA, USA). In MatLab the Fieldtrip ^28^ and the Brainstorm ^29^ toolboxes and custom-written code were used. Two different cortical activation states have been described under urethane anesthesia. ^30, 31^ We confined our analysis to periods of spontaneously activated states, because changes in oscillatory network activity have been mostly described in this state. ^24, 25, 32, 33^

The calculation of the power spectral density (PSD) was estimated by the Welch method (pwelch, NFFT = 1000 pts (i.e. 1 s), 1s Hanning-window with 50% overlap, frequency resolution of 1 Hz). The calculated PSD was normalized by the average between 80 and 100 Hz, where the spectrum shows no oscillatory activity. Neural interactions across LFP frequencies from different recording sites was estimated by calculating the coherence. Coherence was defined as the normalized cross-spectrum (Nolte et al. 2004). Power and absolute coherence values were averaged over the following frequency ranges: theta (4-8 Hz), alpha (9-12 Hz), beta (13–30 Hz) and gamma (55-80 Hz). Data were compared with group-averaged controls.

Because the peaks of oscillatory activity in the power and coherence analysis had non-overlapping maxima between recording sites and animals, we identified individual peaks according published criteria ^24, 34^ and superimposed them for a peak-centered analysis (+/-5 Hz in the beta and +/- 7Hz in the gamma range). Additionally, the oscillatory component was isolated by subtracting pink noise using the FoooF toolbox.32 in Brainstorm ^29^.

### 1.7 Histology and immunochemistry

At the end of the recordings, the animals were sacrificed. Rats were transcardially perfused with 250 ml of 0.1 M phosphate buffered saline (PBS, pH = 7.4), followed by 250 ml of 4% paraformaldehyde in 0.1 M PBS. Brains were removed and postfixed in PFA for 24 h. For cryoprotection, brains were immersed in 30% sucrose solutions and then stored at −80 °C until sectioning. Coronal 40 μm sections were cut with a cryotome (Leica, Germany) and stained with cresyl violet. Electrode trajectories and the placement of the osmotic pumps were verified using a microscope (Zeiss Microscope, Carl Zeiss Microscopy Deutschland GmbH, Germany).

### 1.8 Statistical analysis

Statistics were calculated using GraphPad Prism 10 (GraphPad Software, CA, USA). Statistical significance was assessed using unpaired two-tailed t-tests or one-way ANOVA depending on the number of groups compared. Post hoc tests were calculated only when the omnibus F-values showed significant results. The significance level of all tests was defined to be alpha < 0.05 (* < 0.05; ** < 0.01; *** < 0.001; **** <0.0001).

## Results

### Cerebellar ouabain infusion results in severe dystonia, high frequency gamma and reduced beta oscillations in the networks

The CB-group was infused into the CB with ouabain until the animals (n=8) reached a maximum score on the dystonia rating scale or for a maximum of 72 hours. The mean score achieved by the animals was 3.5 out of 4 (indicating severe sustained dystonia) after an average infusion time of 55 hours. Subsequently, we conducted simultaneous multisite in vivo electrophysiological recordings of local field potentials under urethane anesthesia from the deep cerebellar nuclei (DCN, representing the main output of the cerebellum), the substantia nigra reticulata (SNr, representing the main output of the basal ganglia), the centrolateral-thalamus (Cl, the subcortical anterograde relay nucleus for information flow from the CB to the BG) and the primary motor cortex (M1, Fig. 1A). The CB-group was compared to an untreated control group of animals that were recorded at the same sites in the brain. A power spectral density analysis of the recordings of individual animals from the CB-group revealed the presence of distinct peaks of oscillatory gamma activity at all recording sites (Fig. 3). The mean peak frequency was 65.75 Hz (Fig. 3A). The average high gamma activity (55-80 Hz) was significantly elevated only in the CB-group at the DCN and Cl compared to controls, but not at the other sites. This was likely due to the frequency variation of peaks that cancelled each other out when calculating the average across animals in the gamma band. Consequently, a peak-centered analysis was conducted, whereby power spectral density graphs from individual animals were realigned based on their maximal peaks (+/-7 Hz), and the non-oscillatory 1/f-component was subtracted (using the FOOF toolbox). Peak centered high gamma activity was significantly elevated at all recording sites compared to controls at the same average frequency (Fig. 5A). In accordance with these findings, coherences between the recorded network nodes were significantly elevated in the high gamma range (55-80 Hz) for the CB-group compared to controls for the combinations DCN- SNr, DCN-Cl, M1-Cl and DCN-M1, but not for Cl-SNr and M1-SNr (Suppl. Fig. 1). The frequencies of gamma peaks varied between different animals and combinations and was on average at 67 Hz (Fig. 5B). A peak-centered analysis showed significantly elevated peak centered gamma coherences for all combinations (Suppl. Fig. 4A).

The power spectral density analysis showed significantly reduced beta activity (13-35 Hz) at the Cl and DCN of the CB-group compared to controls (Fig. 2). Beta-coherences were also significantly reduced in most combinations of network nuclei (Suppl. Fig. 1). In the other frequency bands (alpha, theta) no change in average activity levels were found.

**Fig.2:**
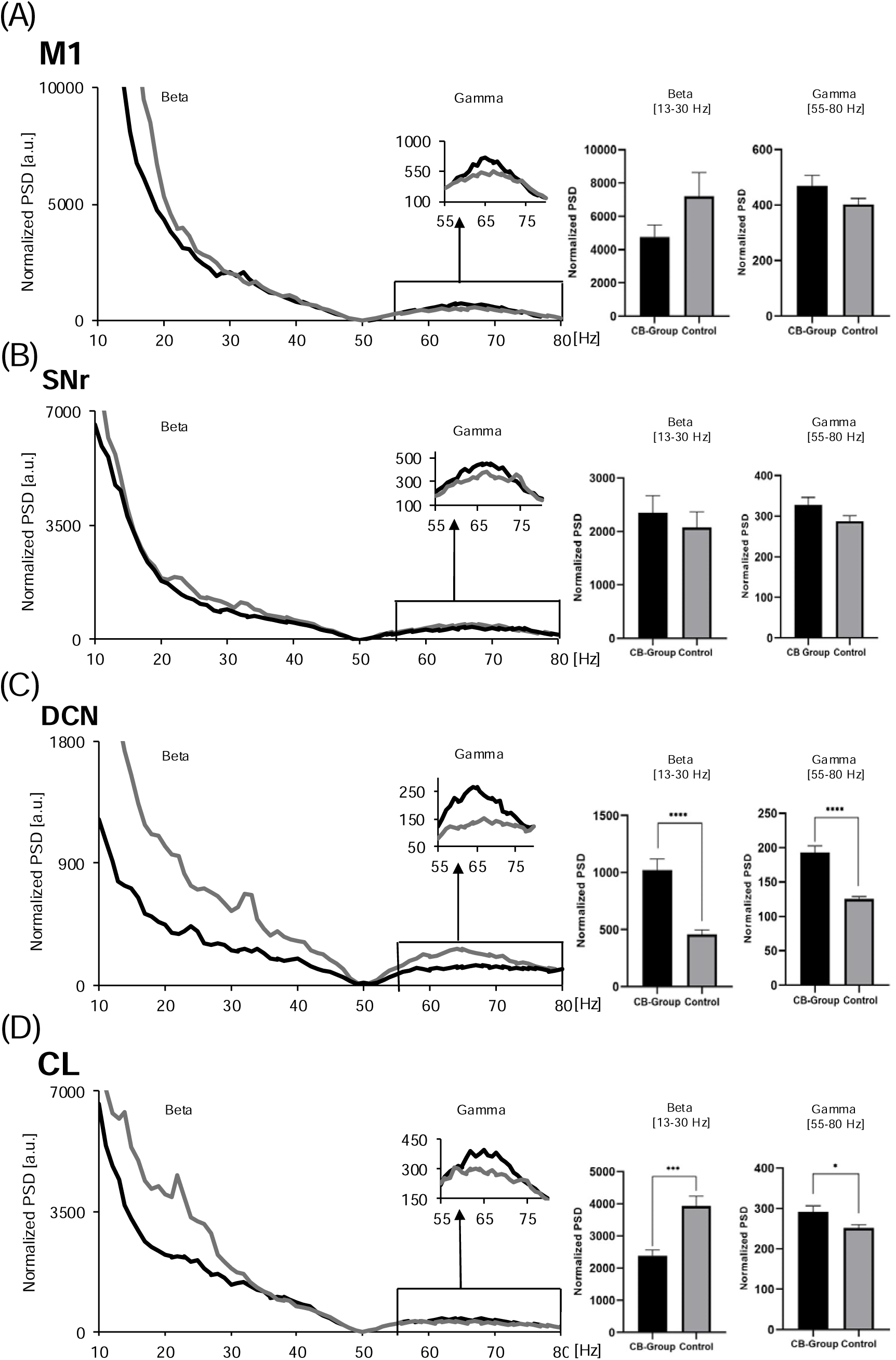
Power spectral density analysis of the CB-Group. **(A)** M1, **(B)** SNr, **(C)** DCN, **(D)** Cl. On the left power plots are shown with insets of magnified segments. On the right bar plots comparing mean beta and gamma power.

**Fig.3:**
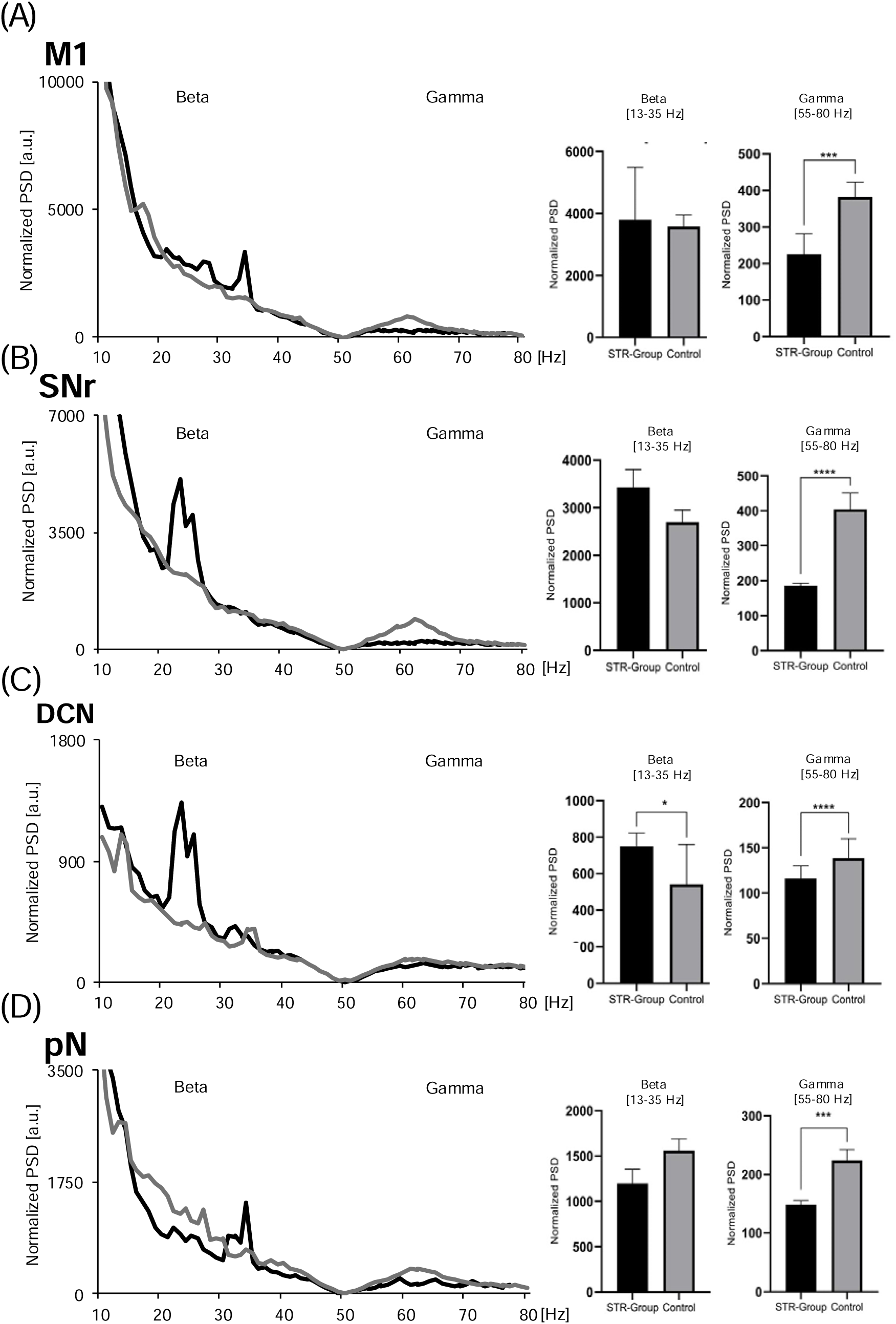
Power spectral density analysis of the STR-Group. **(A)** M1, **(B)** SNr, **(C)** DCN, **(D)** pN. On the left power plots are shown with insets of magnified segments. On the right bar plots comparing mean beta and gamma power.

### Striatal ouabain infusion results in pronounced parkinsonism and elevated network beta activity

Ouabain was infused bilaterally into the striatum (STR-group) until the animals (n=8) reached a maximum score on the Locomotion Disability Scale (LDS) or up to a maximum of 72 hours. The mean LDS score for the animals was 2/3, with an average infusion time of 47.5 hours. As with the electrophysiological experiments conducted on the CB group, the STR group was simultaneously recorded from M1, CB, SNr, and in addition from the pontine nuclei (pN), which serve as the anterograde relay nucleus for information flow from the BG to the CB (Fig. 1B). A power spectral density analysis revealed the presence of significant peaks of oscillatory beta activity in the STR-group with an average frequency of 21.5 Hz (range 20 - 23 Hz), which was not observed in controls (Fig. 3). However, average beta activity (13-35 Hz) was significantly increased only at the SNr (Fig. 3). A peak centered analysis of beta activity (peak +/- 5 Hz) demonstrated significant elevations in beta power at all recorded sites except for the pN compared to controls (Fig. 5).

Mean beta coherence was elevated in the STR-group for the combination M1-SNr, DCN-SNr, pN-M1 and was significantly reduced for pN-SNr and pN-DCN (Suppl. Fig. 2).

Gamma power and gamma coherences were reduced in the STR-group compared to controls and all recording sites and for all coherence combinations (Suppl. Fig. 2).

### Combined striatal and cerebellar ouabain infusion results in dystonia-parkinsonism and elevated network beta activity

Ouabain was infused bilaterally into the STR and the CB (STRCB-group) until the rats (n=8) reached a maximum score on the DRS, or the LDS or until a maximal time of 72 hours had elapsed. The mean DRS score was 3.4/4 and the mean LDS score was 2.8/3, with an average infusion time of 33.5 hours. Electrophysiological recordings were conducted on the M1, SNr, and DCN, and were compared to controls. The animals in the STRCB-group exhibited notable peaks in the beta range, but not in the gamma range (Fig. 4, 5). The beta peaks were distributed between 26 and 31 Hz had an average peak frequency of 28 Hz (Fig. 5). Average beta power (13-35 Hz) was significantly elevated only in the DCN (Fig. 4). To clarify this discrepancy between the visible peaks of oscillatory beta activity and the average power, a peak-centered power analysis was performed, which revealed significant beta power peaks at all recording sites (Fig. 5).

**Fig. 4:**
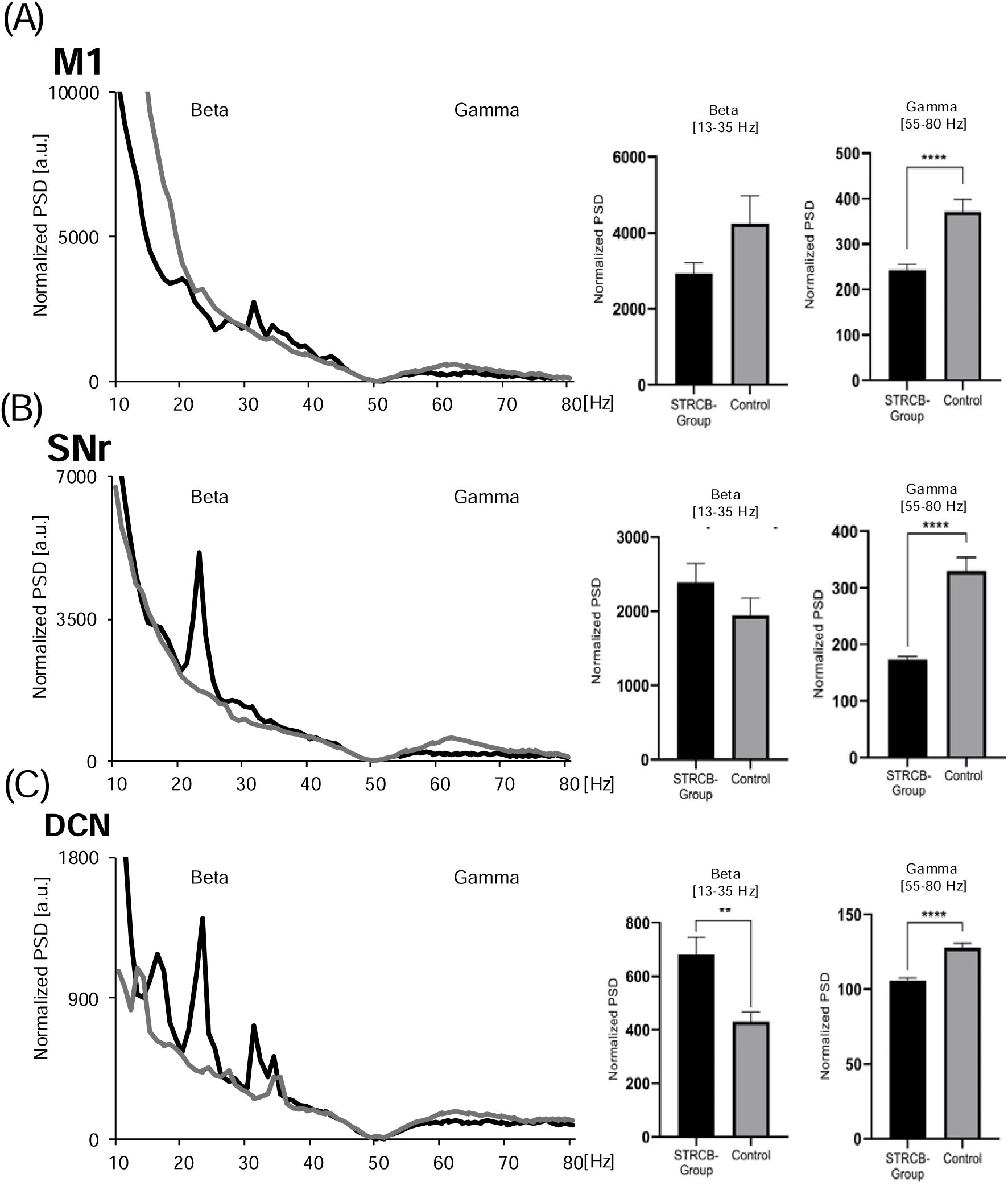
Power spectral density analysis of the STRCB-Group. **(A)** M1, **(B)** SNr, **(C)** DCN, **(D)** cl. On the left power plots are shown with insets of magnified segments. On the right bar plots comparing mean beta and gamma power.

**Fig.5:**
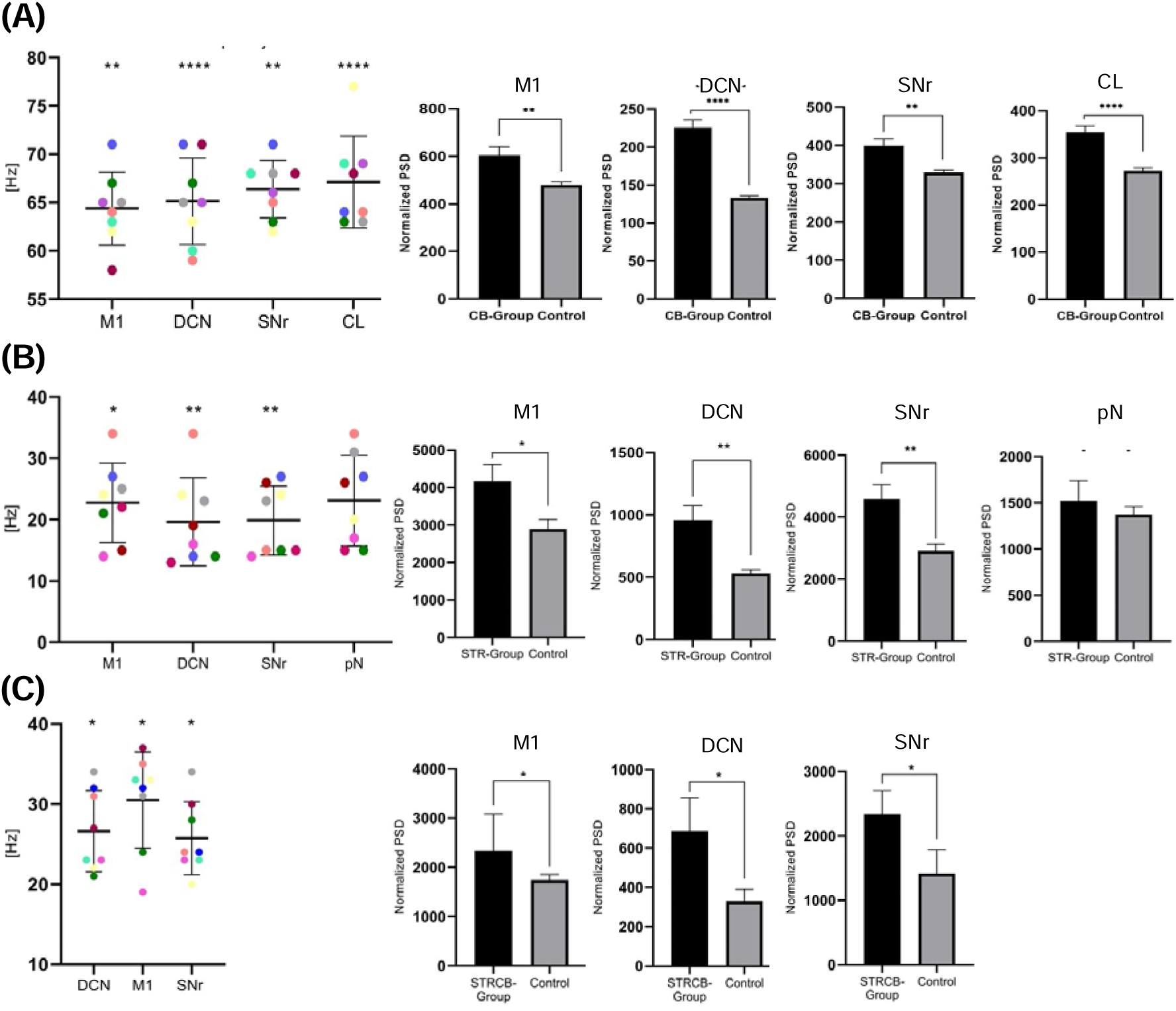
Peak-centered power calculation. **(A)** CB-group. **(B)** STR-group. **(C)** STRCB-group. On the left distribution of peak frequencies are shown. On the right bar plots for the peak- centered power calculation.

Beta coherences were significantly increased only for the combination DCN-SNr in the calculations of average beta and in the peak centered analysis (Suppl. Fig. 3, 4).

In addition, there was a clear reduction in the STRCB-group in average gamma power and average gamma coherence at all recording sites and for the possible combinations (Fig. 4, Suppl. Fig. 3).

## Discussion

There is increasing evidence that a considerable number of movement disorders are characterized by a combined dysfunction of the basal ganglia and the cerebellum.^16, 35, 36^ Nevertheless, for a variety of reasons, research projects have typically concentrated on one or the other of these motor control loops with minimal attention directed towards the other and their interaction. A prime example of a disease for which a combined involvement of the cerebellum and the basal ganglia has been demonstrated is RDP.

In the present study, we conducted a network-level analysis of the interaction between the basal ganglia and the cerebellum in the ouabain model of RDP. This model is ideally suited to dissect this interaction between the BG and the CB, because dystonia and parkinsonism can be selectively induced by infusion of the toxin into the CB or the STR, respectively. Furthermore, evidence indicates that cerebellar-induced dystonia also depends on a functional subcortical interaction between the CB and the BG, suggesting a combined dysfunction of the two motor control loops.^8^ If the observed parkinsonism in RDP is dependent on cerebellar dysfunction has not been investigated, yet.

Our study is the first to demonstrate that symptom-associated increased oscillatory activity originating in the CB or BG is transmitted to the other control loop and M1. This finding is likely to be of great importance for the pathophysiology of the disease, as previous experimental evidence has shown that interrupting the connection between BG and CB can prevent the development of symptoms.^8^

Our findings demonstrate that ouabain infusion into the CB lead to dystonia and results in the generation of high-frequency narrow-bandwidth gamma oscillations, which are subsequently propagated to the SNr and M1. Cerebellar dysfunction has been frequently linked to dystonia in humans (as evidenced by imaging studies in hereditary dystonia and case series in spinocerebellar ataxias) and in animal models of disease before.^35, 37–42^ The question of how the BG and the CB contribute to different dystonic disorders remains a topic of ongoing debate.^35, 39, 42, 43^ While increased Gamma oscillations have been identified in association with hyperkinetic movement disorders such as dystonia and levodopa-induced dyskinesia, recent studies on dystonic patients also demonstrated alterations in lower frequencies.^44–48^ The observed differences between theta and gamma oscillations may be attributed to variations in the brain region recorded or to the heterogeneity in the pathophysiology of patients with dystonia.

In the second part of our study, we demonstrated that bilateral striatal ouabain infusion, which resulted in parkinsonism, led to increased oscillatory beta activity, which was transmitted to the cerebellum and M1. An increase in oscillatory beta activity within the cortico-basal ganglia loop has been strongly linked to akinesia in PD patients as well as in experimental animal models in the past.^24, 25, 49–51^ It seems probable that the increased beta activity and akinesia observed in our RDP model were also induced by changes in striatal dopamine metabolism, as recently described.^11^ It has yet to be determined whether PD-related network changes in the form of enhanced beta oscillations extend to the cerebellum. Nevertheless, it is well-established that the cerebellum plays a pivotal role in tremor-dominant Parkinson’s disease. The dimmer-switch hypothesis of parkinsonian tremor posits that the BG activate tremor, while the cerebello-thalamo-cortical circuit modulates tremor amplitude.^16^

We found increased beta but not gamma activity when infusing ouabain into the striatae and the cerebellum, as would have been expected from the experiments with a single infusion site. It can be speculated that the underlying cellular mechanisms of highly synchronized oscillatory beta and gamma activity are mutually exclusive and that a network can only be dominated by one or the other.

The translational validity of the animal model utilized in the present study is a limitation, particularly when compared to published genetic models and human patients. Nevertheless, the ouabain model offers the unique opportunity to examine the role of the cerebellum and the basal ganglia in the pathophysiology of the disease independently and to characterize their interaction. Additionally, due to spatial constraints on the animal’s head, we were only able to record from a discrete number of network sites within CB and BG. To address this limitation, we selected to record the output nodes of the basal ganglia and cerebellum, as it is probable that their activity reflects the oscillatory activity of the entire loop. Our data were recorded in anesthetized animals; however, as we and others have demonstrated, the activated cortical network states utilized for the analysis closely replicates findings made in awake subjects.^24, 27, 32, 33, 52^

In conclusion, in our study on RDP we are the first to demonstrate that dystonia associated gamma oscillations are transmitted from the CB to the BG and that parkinsonism associated beta oscillations get transmitted from the BG to the CB. We hypothesize that this mutual influence via oscillatory synchronization is of great importance for the pathophysiology of RDP and the generation of symptoms. The extent to which a combined dysfunction of the CB and BG is also relevant in other movement disorders should be investigated in future research. A more detailed understanding of the coordination of the BG and the CB in movement disorders could facilitate the development of targeted, circuit-specific neuromodulatory therapies, such as deep brain stimulation.

## Documentation of Author roles

1. Research Project: A. Conception, B. Organization, C. Execution
2. Statistical Analysis: A. Design, B. Execution, C. Review and Critique
3. Manuscript preparation: A. Writing of the first draft, B. Review and Critique

M.M. 1B, 1C, 2A, 2B, 3A, 3B

J.A.N..: 1B, 1C, 3B

V.V.N.: 2C, 3B

C.v.R.: 1A, 1B, 1C, 2A, 2C, 3A, 3B

## Financial disclosure of all authors (for the preceding 12 month)

M.M., J.A.N., V.V.N., C.v.R. none.

## Supplementary Figure Legends

**Suppl. Fig.1.:**
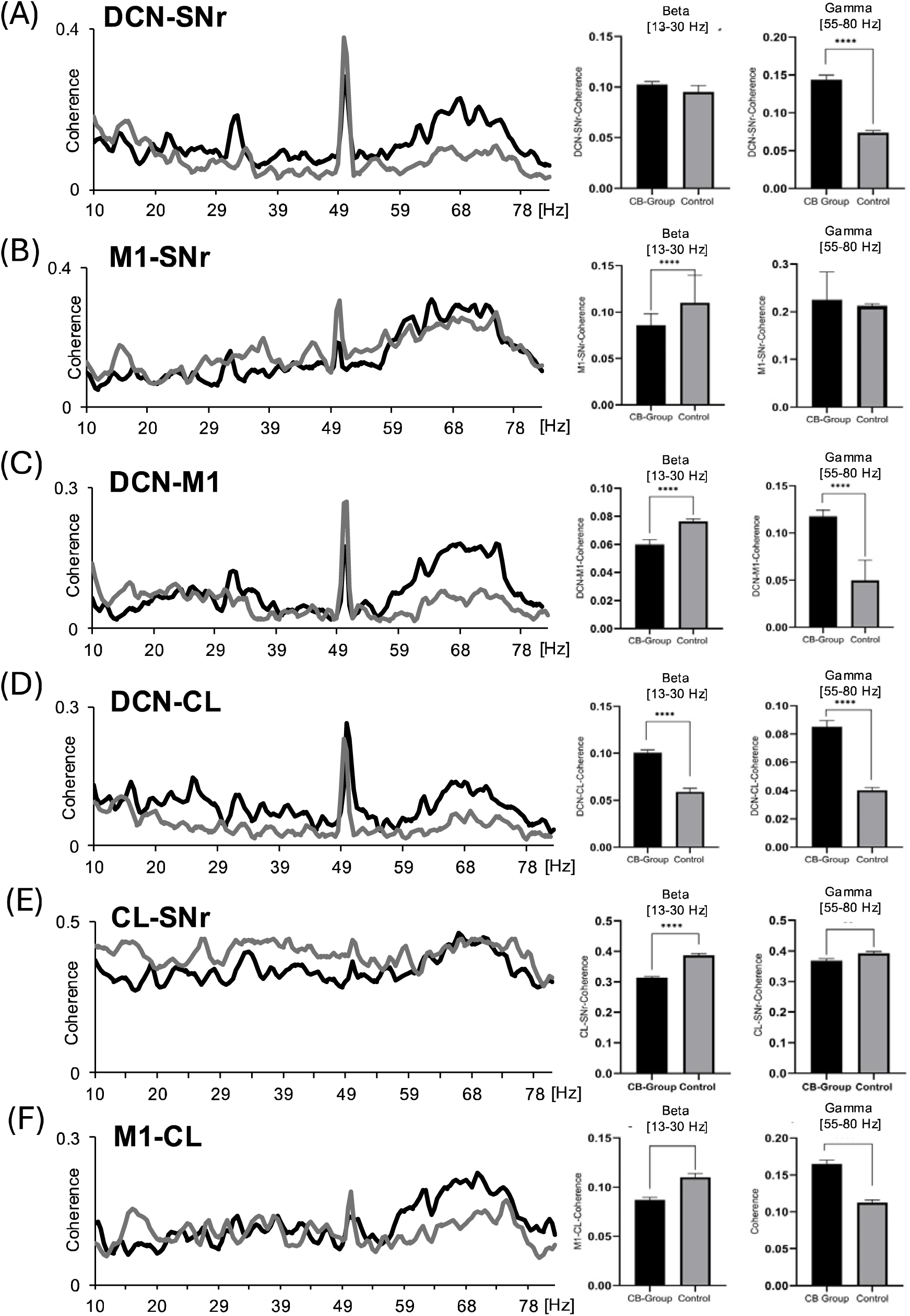
Coherence analysis of the CB-Group. **(A)** DCN-SNr, **(B)** DCN-Cl, **(C)** Cl-SNR, **(D)** M1-Cl, (E) M1-SNr, (F) DCN-M1. On the left coherence spectra are shown. On the right bar plots comparing mean beta and gamma coherences.

**Suppl. Fig.2:**
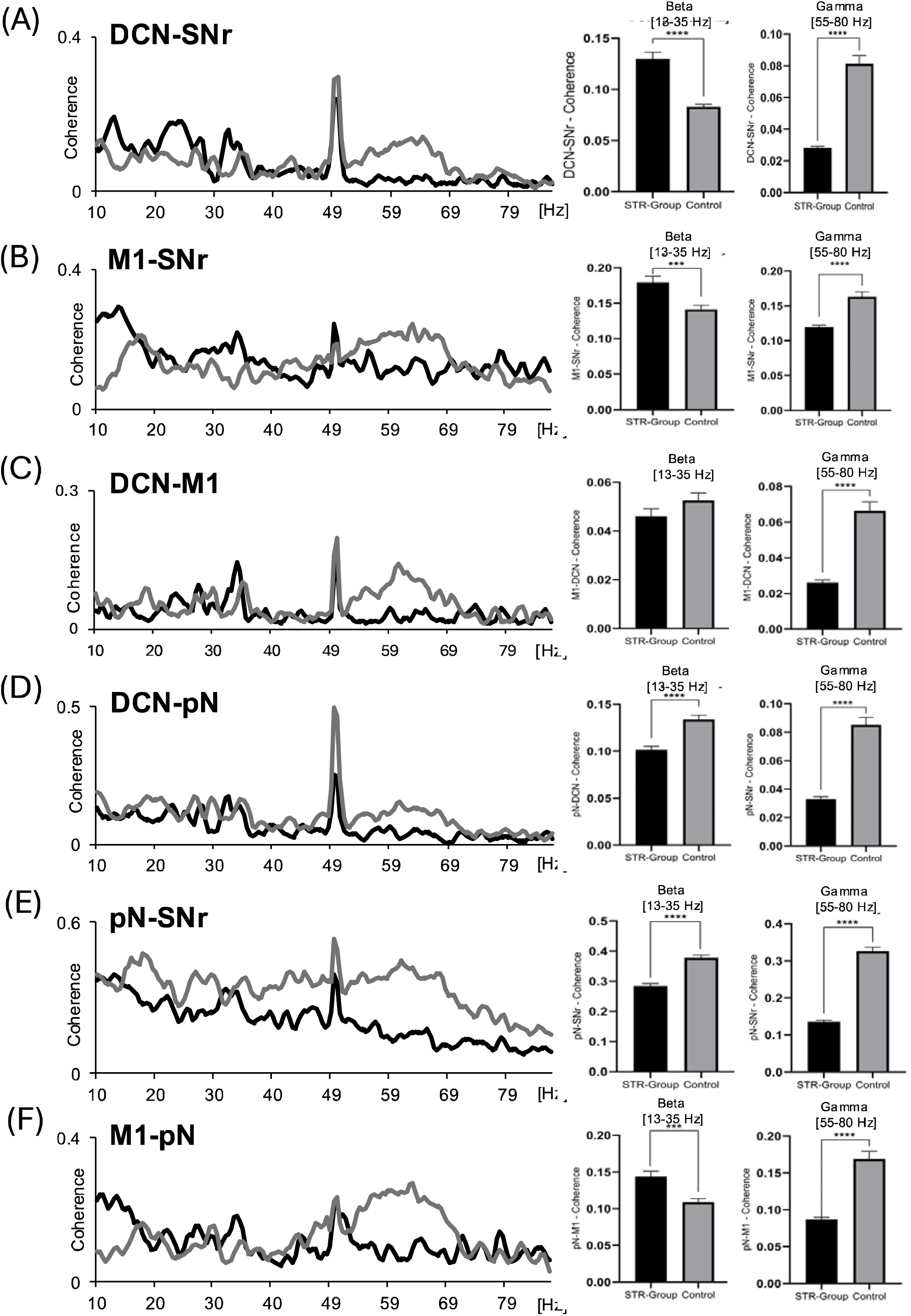
Coherence analysis of the STR-Group. **(A)** M1-DCN, **(B)** M1-SNr, **(C)** DCN- SNr, **(D)** pN-M1, (E) pN-SNr, (F) pN-DCN. On the left coherence spectra are shown. On the right bar plots comparing mean beta and gamma coherences.

**Suppl. Fig.3:**
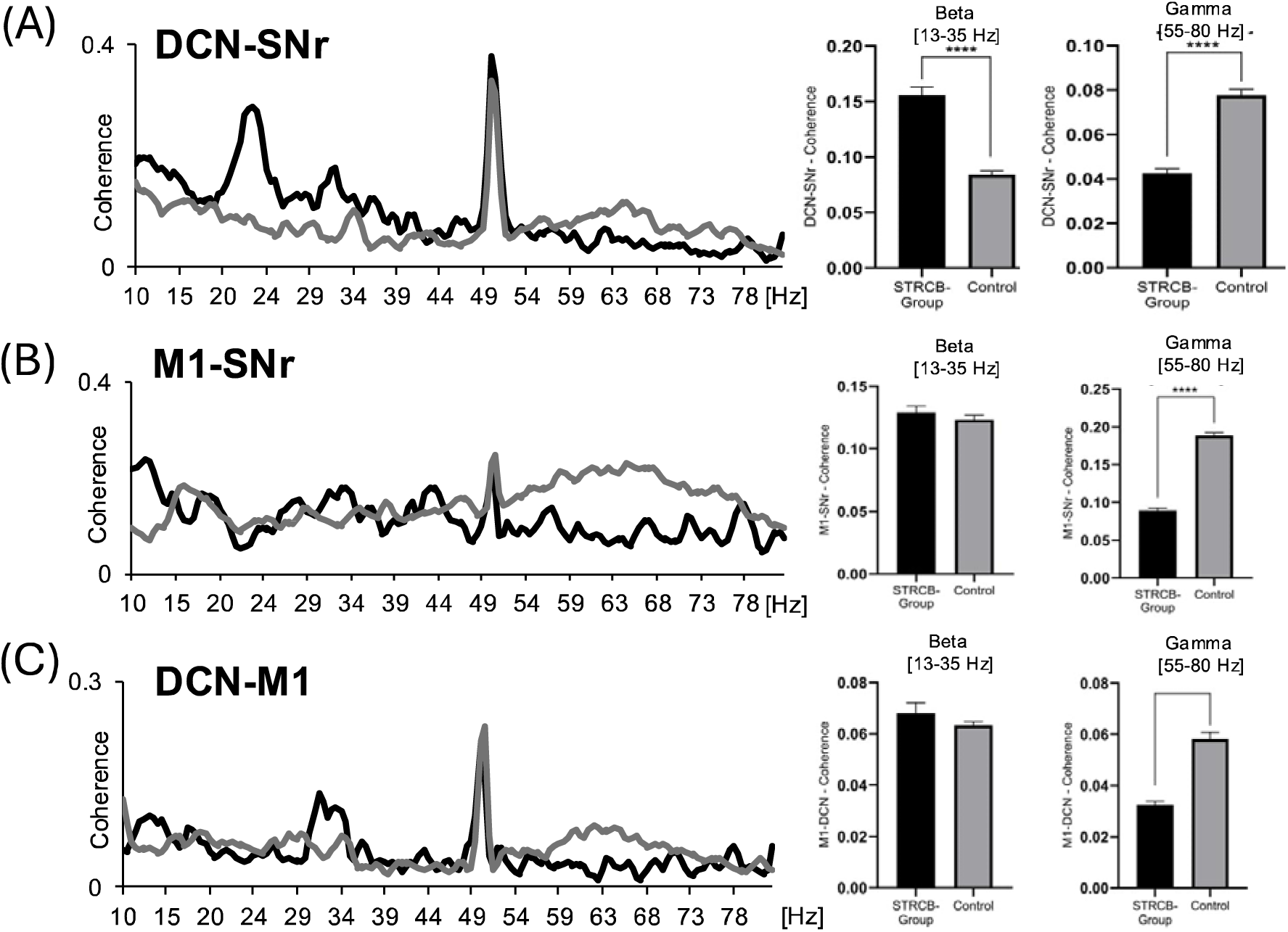
Coherence analysis of the STRCB-Group. **(A)** DCN-SNr, **(B)** M1-SNr, **(C)** M1- DCN. On the left coherence plots are shown. On the right bar plots comparing mean beta and gamma coherences.

**Suppl. Fig.4:**
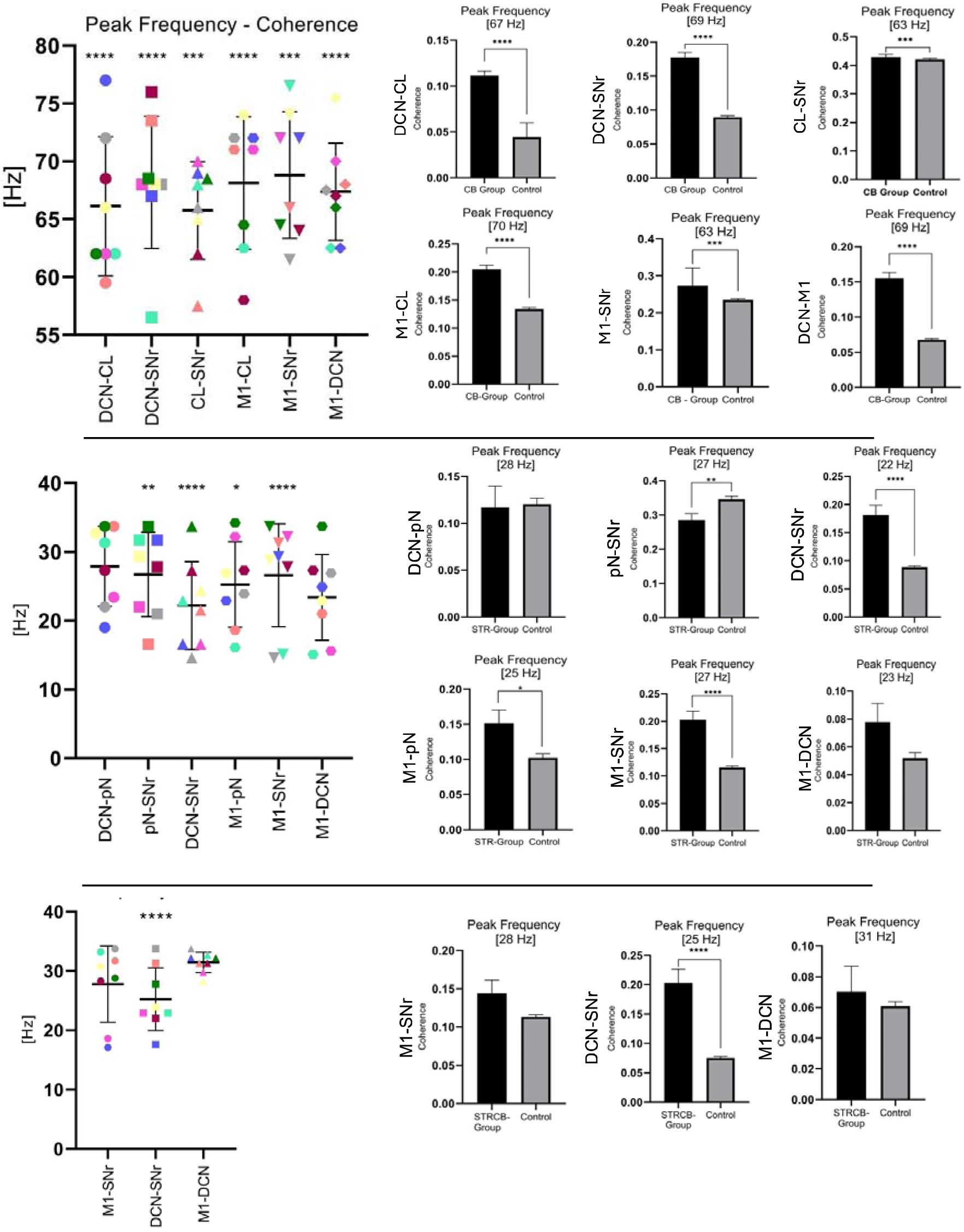
Peak-centered coherence calculation. **(A)** CB-group. **(B)** STR-group. **(C)** STRCB-group. On the left distribution of peak frequencies are shown. On the right bar plots for the peak-centered coherence calculation.

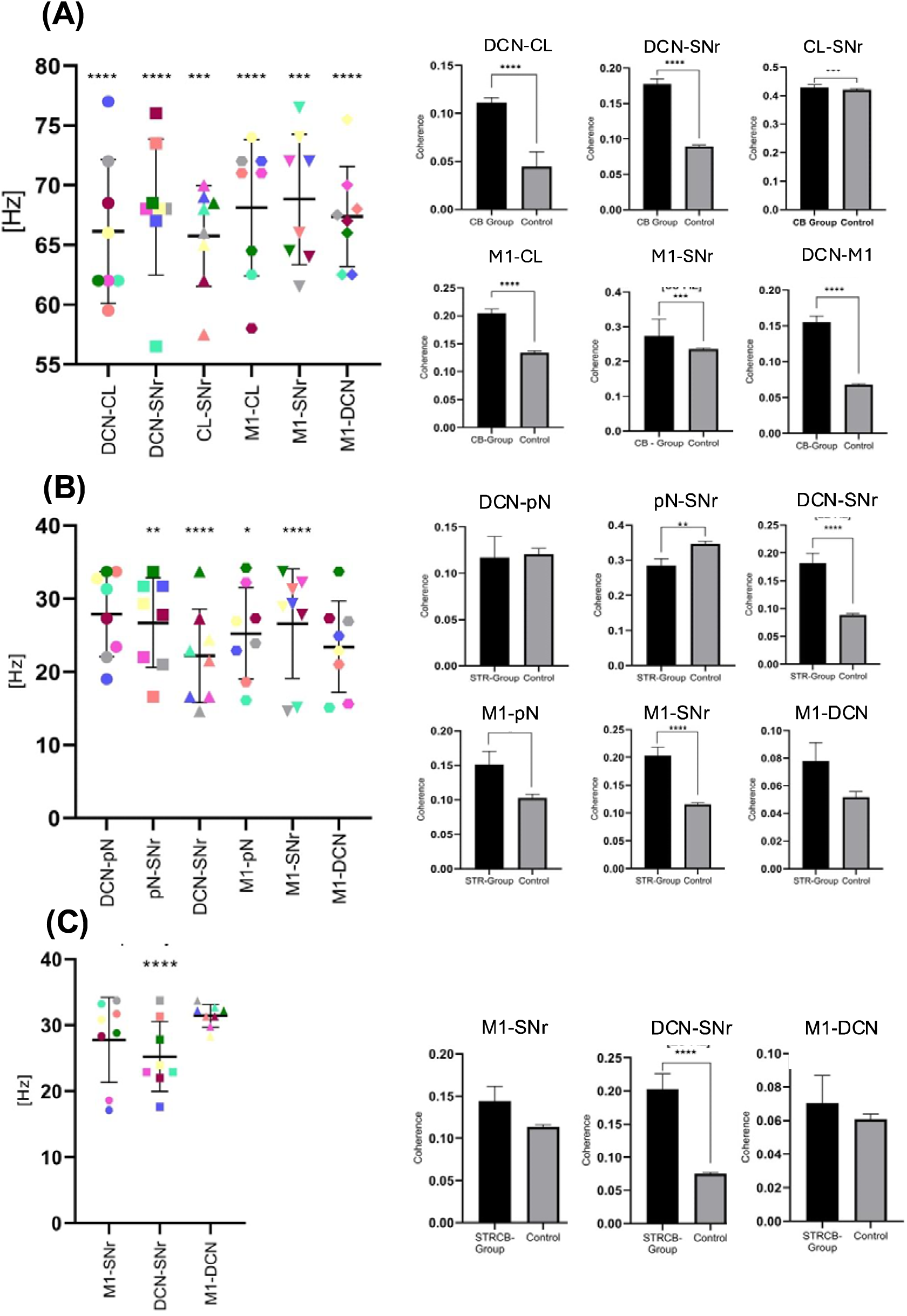

**Suppl. Tabl. 1.**

| **Coordinates of osmotic pump** | **Cerebellum** | **Striatum** |
| --- | --- | --- |
| **AP** | - 12.4 | - 1.8 |
| **ML** | +/- 0.0 | +4.2/ -4.2 |
| **DV** | - 5.0 | - 5.1 |

**Suppl. Tabl. 2.**

| **Coordinates of microelectrodes** | **DCN** | **SNr** | **CL** | **pN** | **M1 (ECoG)** |
| --- | --- | --- | --- | --- | --- |
| **AP** | - 11.3 | - 4.8 | -3.3 | -6.8 | + 2.7 |
| **ML** | + 3.2 | -2.5 | -1.4 | +0.5 | + 2.0 |
| **DV** | -5.8 to -6.1 | - 7.6 to -8.6 | -4.9 to -5.5 | - 9.4 to - 10.3 | / |

## Notes

### Competing Interest Statement

The authors have declared no competing interest.

## References

1. Deutschlander A, Asmus F, Gasser T, Steude U, Botzel K. Sporadic rapid-onset dystonia-parkinsonism syndrome: failure of bilateral pallidal stimulation. Mov Disord 2005;20(2):254–257.

2. Kamm C, Fogel W, Wachter T, et al. Novel ATP1A3 mutation in a sporadic RDP patient with minimal benefit from deep brain stimulation. Neurology 2008;70(16 Pt 2):1501–1503.

3. de Carvalho Aguiar P, Sweadner KJ, Penniston JT, et al. Mutations in the Na+/K+ - ATPase alpha3 gene ATP1A3 are associated with rapid-onset dystonia parkinsonism. Neuron 2004;43(2):169–175.

4. Kaplan JH. Biochemistry of Na,K-ATPase. Annu Rev Biochem 2002;71:511–535.

5. Bottger P, Tracz Z, Heuck A, Nissen P, Romero-Ramos M, Lykke-Hartmann K. Distribution of Na/K-ATPase alpha 3 isoform, a sodium-potassium P-type pump associated with rapid-onset of dystonia parkinsonism (RDP) in the adult mouse brain. J Comp Neurol 2011;519(2):376–404.

6. Einholm AP, Toustrup-Jensen MS, Holm R, Andersen JP, Vilsen B. The rapid-onset dystonia parkinsonism mutation D923N of the Na+, K+-ATPase alpha3 isoform disrupts Na+ interaction at the third Na+ site. J Biol Chem 2010;285(34):26245–26254.

7. Holm TH, Lykke-Hartmann K. Insights into the Pathology of the alpha3 Na(+)/K(+)- ATPase Ion Pump in Neurological Disorders; Lessons from Animal Models. Front Physiol 2016;7:209.

8. Calderon DP, Fremont R, Kraenzlin F, Khodakhah K. The neural substrates of rapid-onset Dystonia-Parkinsonism. Nat Neurosci 2011;14(3):357–365.

9. Chen CH, Fremont R, Arteaga-Bracho EE, Khodakhah K. Short latency cerebellar modulation of the basal ganglia. Nat Neurosci 2014;17(12):1767–1775.

10. Fremont R, Calderon DP, Maleki S, Khodakhah K. Abnormal high-frequency burst firing of cerebellar neurons in rapid-onset dystonia-parkinsonism. J Neurosci 2014;34(35):11723–11732.

11. Rauschenberger L, Knorr S, Al-Zuraiqi Y, Tovote P, Volkmann J, Ip CW. Striatal dopaminergic dysregulation and dystonia-like movements induced by sensorimotor stress in a pharmacological mouse model of rapid-onset dystonia-parkinsonism. Experimental neurology 2020;323:113109.

12. Akkal D, Dum RP, Strick PL. Supplementary motor area and presupplementary motor area: targets of basal ganglia and cerebellar output. J Neurosci 2007;27(40):10659–10673.

13. Bostan AC, Dum RP, Strick PL. The basal ganglia communicate with the cerebellum. Proc Natl Acad Sci U S A 2010;107(18):8452–8456.

14. Clower DM, Dum RP, Strick PL. Basal ganglia and cerebellar inputs to ‘AIP’. Cereb Cortex 2005;15(7):913–920.

15. Hoover JE, Strick PL. The organization of cerebellar and basal ganglia outputs to primary motor cortex as revealed by retrograde transneuronal transport of herpes simplex virus type 1. J Neurosci 1999;19(4):1446–1463.

16. Helmich RC. The cerebral basis of Parkinsonian tremor: A network perspective. Mov Disord 2018;33(2):219–231.

17. Bhuvanasundaram R, Krzyspiak J, Khodakhah K. Subthalamic Nucleus Modulation of the Pontine Nuclei and Its Targeting of the Cerebellar Cortex. J Neurosci 2022;42(28):5538–5551.

18. Chen CC, Lin WY, Chan HL, et al. Stimulation of the subthalamic region at 20 Hz slows the development of grip force in Parkinson’s disease. Experimental neurology 2011;231(1):91–96.

19. Hoshi E, Tremblay L, Feger J, Carras PL, Strick PL. The cerebellum communicates with the basal ganglia. Nat Neurosci 2005;8(11):1491–1493.

20. Ichinohe N, Mori F, Shoumura K. A di-synaptic projection from the lateral cerebellar nucleus to the laterodorsal part of the striatum via the central lateral nucleus of the thalamus in the rat. Brain Res 2000;880(1-2):191–197.

21. Buzsaki G, Anastassiou CA, Koch C. The origin of extracellular fields and currents-- EEG, ECoG, LFP and spikes. Nat Rev Neurosci 2012;13(6):407–420.

22. Friston KJ, Bastos AM, Pinotsis D, Litvak V. LFP and oscillations-what do they tell us? Current opinion in neurobiology 2015;31:1–6.

23. Paxinos G, Watson C. The rat brain in stereotaxix coordinates 7th edition. Academic Press 2013.

24. Beck MH, Haumesser JK, Kuhn J, Altschuler J, Kuhn AA, van Riesen C. Short- and long-term dopamine depletion causes enhanced beta oscillations in the cortico-basal ganglia loop of parkinsonian rats. Experimental neurology 2016;286:124–136.

25. Haumesser JK, Beck MH, Pellegrini F, et al. Subthalamic beta oscillations correlate with dopaminergic degeneration in experimental parkinsonism. Experimental neurology 2021;335:113513.

26. Haumesser JK, Kuhn J, Guttler C, et al. Acute In Vivo Electrophysiological Recordings of Local Field Potentials and Multi-unit Activity from the Hyperdirect Pathway in Anesthetized Rats. J Vis Exp 2017(124).

27. Kuhn J, Haumesser JK, Beck MH, et al. Differential effects of levodopa and apomorphine on neuronal population oscillations in the cortico-basal ganglia loop circuit in vivo in experimental parkinsonism. Experimental neurology 2017;298(Pt A):122–133.

28. Oostenveld R, Fries P, Maris E, Schoffelen JM. FieldTrip: Open source software for advanced analysis of MEG, EEG, and invasive electrophysiological data. Comput Intell Neurosci 2011;2011:156869.

29. Tadel F, Baillet S, Mosher JC, Pantazis D, Leahy RM. Brainstorm: a user-friendly application for MEG/EEG analysis. Comput Intell Neurosci 2011;2011:879716.

30. Magill PJ, Bolam JP, Bevan MD. Relationship of activity in the subthalamic nucleus-globus pallidus network to cortical electroencephalogram. J Neurosci 2000;20(2):820–833.

31. Steriade M. Corticothalamic resonance, states of vigilance and mentation. Neuroscience 2000;101(2):243–276.

32. Mallet N, Pogosyan A, Marton LF, Bolam JP, Brown P, Magill PJ. Parkinsonian beta oscillations in the external globus pallidus and their relationship with subthalamic nucleus activity. J Neurosci 2008;28(52):14245–14258.

33. Mallet N, Pogosyan A, Sharott A, et al. Disrupted dopamine transmission and the emergence of exaggerated beta oscillations in subthalamic nucleus and cerebral cortex. J Neurosci 2008;28(18):4795–4806.

34. Delaville C, McCoy AJ, Gerber CM, Cruz AV, Walters JR. Subthalamic nucleus activity in the awake hemiparkinsonian rat: relationships with motor and cognitive networks. J Neurosci 2015;35(17):6918–6930.

35. Prudente CN, Hess EJ, Jinnah HA. Dystonia as a network disorder: what is the role of the cerebellum? Neuroscience 2014;260:23–35.

36. Tewari A, Fremont R, Khodakhah K. It’s not just the basal ganglia: Cerebellum as a target for dystonia therapeutics. Mov Disord 2017;32(11):1537–1545.

37. Fremont R, Tewari A, Angueyra C, Khodakhah K. A role for cerebellum in the hereditary dystonia DYT1. Elife 2017;6.

38. Fremont R, Tewari A, Khodakhah K. Aberrant Purkinje cell activity is the cause of dystonia in a shRNA-based mouse model of Rapid Onset Dystonia-Parkinsonism. Neurobiol Dis 2015;82:200–212.

39. Jinnah HA, Hess EJ. A new twist on the anatomy of dystonia: the basal ganglia and the cerebellum? Neurology 2006;67(10):1740–1741.

40. Mariotti C, Alpini D, Fancellu R, et al. Spinocerebellar ataxia type 17 (SCA17): oculomotor phenotype and clinical characterization of 15 Italian patients. J Neurol 2007;254(11):1538–1546.

41. Sethi KD, Jankovic J. Dystonia in spinocerebellar ataxia type 6. Mov Disord 2002;17(1):150–153.

42. Zoons E, Booij J, Nederveen AJ, Dijk JM, Tijssen MA. Structural, functional and molecular imaging of the brain in primary focal dystonia--a review. Neuroimage 2011;56(3):1011–1020.

43. Neychev VK, Gross RE, Lehericy S, Hess EJ, Jinnah HA. The functional neuroanatomy of dystonia. Neurobiol Dis 2011;42(2):185–201.

44. Miocinovic S, Swann NC, de Hemptinne C, Miller A, Ostrem JL, Starr PA. Cortical gamma oscillations in isolated dystonia. Parkinsonism Relat Disord 2018;49:104–105.

45. Neumann WJ, Horn A, Ewert S, et al. A localized pallidal physiomarker in cervical dystonia. Ann Neurol 2017;82(6):912–924.

46. Olaru M, Cernera S, Hahn A, et al. Motor network gamma oscillations in chronic home recordings predict dyskinesia in Parkinson’s disease. Brain 2024;147(6):2038–2052.

47. Scheller U, Lofredi R, van Wijk BCM, et al. Pallidal low-frequency activity in dystonia after cessation of long-term deep brain stimulation. Mov Disord 2019;34(11):1734–1739.

48. Wang DD, de Hemptinne C, Miocinovic S, et al. Subthalamic local field potentials in Parkinson’s disease and isolated dystonia: An evaluation of potential biomarkers. Neurobiol Dis 2016;89:213–222.

49. Kuhn AA, Kupsch A, Schneider GH, Brown P. Reduction in subthalamic 8-35 Hz oscillatory activity correlates with clinical improvement in Parkinson’s disease. Eur J Neurosci 2006;23(7):1956–1960.

50. Neumann WJ, Degen K, Schneider GH, et al. Subthalamic synchronized oscillatory activity correlates with motor impairment in patients with Parkinson’s disease. Mov Disord 2016;31(11):1748–1751.

51. Neumann WJ, Staub-Bartelt F, Horn A, et al. Long term correlation of subthalamic beta band activity with motor impairment in patients with Parkinson’s disease. Clin Neurophysiol 2017;128(11):2286–2291.

52. Hammond C, Bergman H, Brown P. Pathological synchronization in Parkinson’s disease: networks, models and treatments. Trends Neurosci 2007;30(7):357–364.

